# Chaperoning the chaperones: Proteomic analysis of the SMN complex reveals conserved and etiologic connections to the proteostasis network

**DOI:** 10.1101/2024.05.15.594402

**Authors:** A. Gregory Matera, Rebecca E. Steiner, C. Alison Mills, Laura E. Herring, Eric L. Garcia

## Abstract

Molecular chaperones and co-chaperones are highly conserved cellular components that perform variety of duties related to the proper three-dimensional folding of the proteome. The web of factors that carries out this essential task is called the proteostasis network (PN). Ribonucleoproteins (RNPs) represent an underexplored area in terms of the connections they make with the PN. The Survival Motor Neuron (SMN) complex is an RNP assembly chaperone and serves as a paradigm for studying how specific small nuclear (sn)RNAs are identified and paired with their client substrate proteins. SMN protein is the eponymous component of a large complex required for the biogenesis of uridine-rich small nuclear ribonucleoproteins (U-snRNPs) and localizes to distinct membraneless organelles in both the nucleus and cytoplasm of animal cells. SMN forms the oligomeric core of this complex, and missense mutations in its YG box self-interaction domain are known to cause Spinal Muscular Atrophy (SMA). The basic framework for understanding how snRNAs are assembled into U-snRNPs is known, the pathways and mechanisms used by cells to regulate their biogenesis are poorly understood. Given the importance of these processes to normal development as well as neurodegenerative disease, we set out to identify and characterize novel SMN binding partners.

Here, we carried out affinity purification mass spectrometry (AP-MS) of SMN using stable fly lines exclusively expressing either wildtype or SMA-causing missense alleles. Bioinformatic analyses of the pulldown data, along with comparisons to proximity labeling studies carried out in human cells, revealed conserved connections to at least two other major chaperone systems including heat shock folding chaperones (HSPs) and histone/nucleosome assembly chaperones. Notably, we found that heat shock cognate protein Hsc70-4 and other HspA family members preferentially interacted with SMA-causing alleles of SMN. Hsc70-4 is particularly interesting because its mRNA is aberrantly sequestered by a mutant form of TDP-43 in mouse and *Drosophila* ALS (Amyotrophic Lateral Sclerosis) disease models. Most important, a missense allele of Hsc70-4 (HspA8 in mammals) was recently identified as a bypass suppressor of the SMA phenotype in mice. Collectively, these findings suggest that chaperone-related dysfunction lies at the etiological root of both ALS and SMA.

## Introduction

Cellular stressors, both extrinsic and intrinsic, come in a countless variety of shapes and flavors but they all lead to the same end: macromolecular chaos. In order to survive the pull of these entropic forces, at least for a lifetime, organisms have evolved defense mechanisms that help maintain homeostasis in a dynamic environment. These defense mechanisms orchestrate the activation of stress response pathways across tissues and organs to promote cellular and, ultimately, organismal health [1]. Heat stress is unsurprisingly among the most significant barriers to life. All organisms respond to the presence of too much heat by inducing the synthesis of heat shock proteins, or HSPs [2]. Indeed HSPs, are highly conserved in all three kingdoms of life and participate in protein quality control, reviewed in [3].

Proteostasis (protein homeostasis) is a term used to describe the overall process of maintaining a functional proteome. The web of cellular components that carries out this essential task is called the proteostasis network, or PN [1]. A natural aspect of proteostasis involves the degradation and/or recycling of proteins and complexes that are misfolded beyond repair, aggregated or have simply reached the end of their normal lifetimes. Thus the PN includes the ubiquitin proteosome system (UPS) along with several membrane-associated processes that we collectively term autophagy [4]. Additionally, the PN also includes the HSPs that serve as molecular chaperones, many of which are constitutively expressed along with their close partners (co-chaperones), reviewed in [5, 6].

Molecular chaperones and co-chaperones not only prevent co-translational misfolding, but also carry out re-folding of stress-denatured proteins, re-directing non-native intermediates to their native states [6, 7]. Thus, a molecular chaperone is often defined as a protein that helps another protein to acquire its active conformation, without actually being present in the final product [8]. Distinct classes of structurally unrelated chaperones exist in cells, forming cooperative pathways and sub-networks [9]. HSPs belong to the broad category of ‘folding’ chaperones, that also includes the ring-shaped chaperonins [7]. The so-called ‘assembly’ chaperones represent another class of molecular chaperones that help carry out the biogenesis of large macromolecular complexes such as nucleosomes, proteasomes, ribosomes and spliceosomes [10–15].

The Survival Motor Neuron (SMN) complex directs assembly of the uridine-rich small nuclear ribonucleoproteins (U-snRNPs) that make up the spliceosome [16, 17]. Because the SMN complex is essential for proper formation of the RNP, but is not part of the final particle [18], it can be viewed as an assembly chaperone (see [13, 19, 20] for details). In human cells, the complex is composed of SMN and eight main partner proteins [13, 19, 21], collectively called Gemins (Gem2-8 plus STRAP/UNRIP. In *Drosophilids* and other Dipteran genomes, the Gem6•7•8 subunit has been lost [22]. The best-studied client substrates of the SMN complex are the Sm-proteins and U-rich snRNAs [13, 17], although it likely plays a role in facilitating assembly of other RNP classes as well [23–25].

The core of the complex is composed of SMN•Gem2 dimers which, *in vitro*, are fully sufficient for catalyzing assembly of the heteroheptameric Sm-ring onto human or fruitfly snRNAs [26]. Higher-order multimerization of SMN•Gem2 dimers is required for metazoan (but not fission yeast) viability [27]. Note that budding yeast genomes have lost SMN and all of the other Gemins, except Gem2 [13]. Given the fact that each of these organisms have introns and express spliceosomal snRNPs, it seems likely that the rest of the Gemins are involved in regulatory or other aspects of SMN function.

Although the basic framework for understanding how RNPs traffic through the cell during their maturation cycle is known and is conserved among metazoans [28, 29], the pathways and mechanisms used by cells to regulate snRNP biogenesis are poorly understood [13]. How are the activities of other macromolecular assembly machineries connected to snRNP biogenesis and how do they change in disease states? What signaling pathways are used to coordinate these essential cellular functions and how are the signals received? To begin to address these questions, we decided to search beyond the Gemins for additional SMN binding partners. Over the past decade, we have developed *Drosophila* as model system to understand SMN function using an allelic series of transgenic flies expressing SMA-causing missense mutations that recapitulates the full range of phenotypic severity seen in human patients [22, 27, 30–37]

Here, we carried out affinity purification, coupled with mass spectrometry (AP-MS), to identify novel SMN binding partners present in embryonic lysates of wildtype and hypomorphic SMA animal models. Bioinformatic analyses of the hits, along with comparisons to proximity labeling data from human cells revealed connections to at least two other major chaperone systems. Links to the proteostasis network of folding chaperones were even more apparent in the lysates of SMA-causing missense alleles. In particular, we identified the heat shock cognate protein Hsc70-4 and other Hsc70/Hsp70 family members. Hsc70-4 is notable because the mRNA encoding this protein is targeted by a dominant negative, ALS (Amyotrophic Lateral Sclerosis) causing form of TDP-43 in mouse and *Drosophila* disease models [38]. Furthermore, a missense allele of Hsc70-4 (HspA8 in mammals) was recently identified as a suppressor of the SMA phenotype in mice [39].

Taken together, these findings leave little doubt that the underlying neuromuscular phenotypes of SMA and ALS stem from defects in a common pathway served by heat shock folding chaperones.

## Results and Discussion

Until recently, SMA was untreatable and recognized as the most prevalent genetic cause of early childhood mortality [40–42]. Humans have two paralogous SMN genes, named *SMN1* and *SMN2*. SMA is typically caused by deletion of both copies of *SMN1*; left on its own, *SMN2* does not provide enough full-length SMN protein to fully compensate for loss of *SMN1* [43, 44]. Notably, a small number of SMA patients present with a deletion in one copy of *SMN1* and a missense mutation in the other copy [42, 45]. Patients bearing zero copies of *SMN2* together with a homozygous missense mutation in *SMN1* are even more exceptional [46]. Thus, disease presentation in humans varies dramatically, depending on the *SMN1* allele, and the number of *SMN2* gene copies present in the background (which can vary from 0 to 6,).

Our use of *Drosophila melanogaster* as a model system to study SMN function solves a number of the problems that confound phenotypic interpretation in humans and mammals. First, human *SMN2* copy number variation can also mask the phenotype of *SMN1* point mutations [27, 47, 48]. Second, alternative splicing of *SMN2* creates a feedback loop [49, 50] that can negatively regulate SMN expression. Third, purification of native or epitope-tagged SMN from cell lines or tissues may limit the number of binding partners to those that are expressed in a given cell lineage. In flies, there is only one *Smn* gene and its protein coding region is located within a single, constitutively expressed exon.

We have developed a set of fly stocks whose survival depends entirely on transgenic expression of Flag-SMN protein [34, 35]. That is, these animals do not express any endogenous SMN (maternally or zygotically). These flies have the overall genotype: *Smn^X7/X7^,Flag-Smn^Tg/Tg^*, where *Smn^X7^* is a null allele and *Tg* denotes a: *WT, D20V, G73R, I93F* or *G210C* transgene (Fig. 1A). Thus, these models arguably provide a more accurate readout of SMN protein function because the mutants can be expressed and analyzed in the absence of wild-type SMN. Furthermore, the stocks are ideal for carrying out proteomics because we do not have to perform crosses to obtain animals of the desired genotype and we are able to collect large quantities of material. We chose to use embryos because they contain a wide-variety of cell types and they naturally express ∼100x the amount of SMN protein present in larval or adult stages [36].

**Figure 1:**
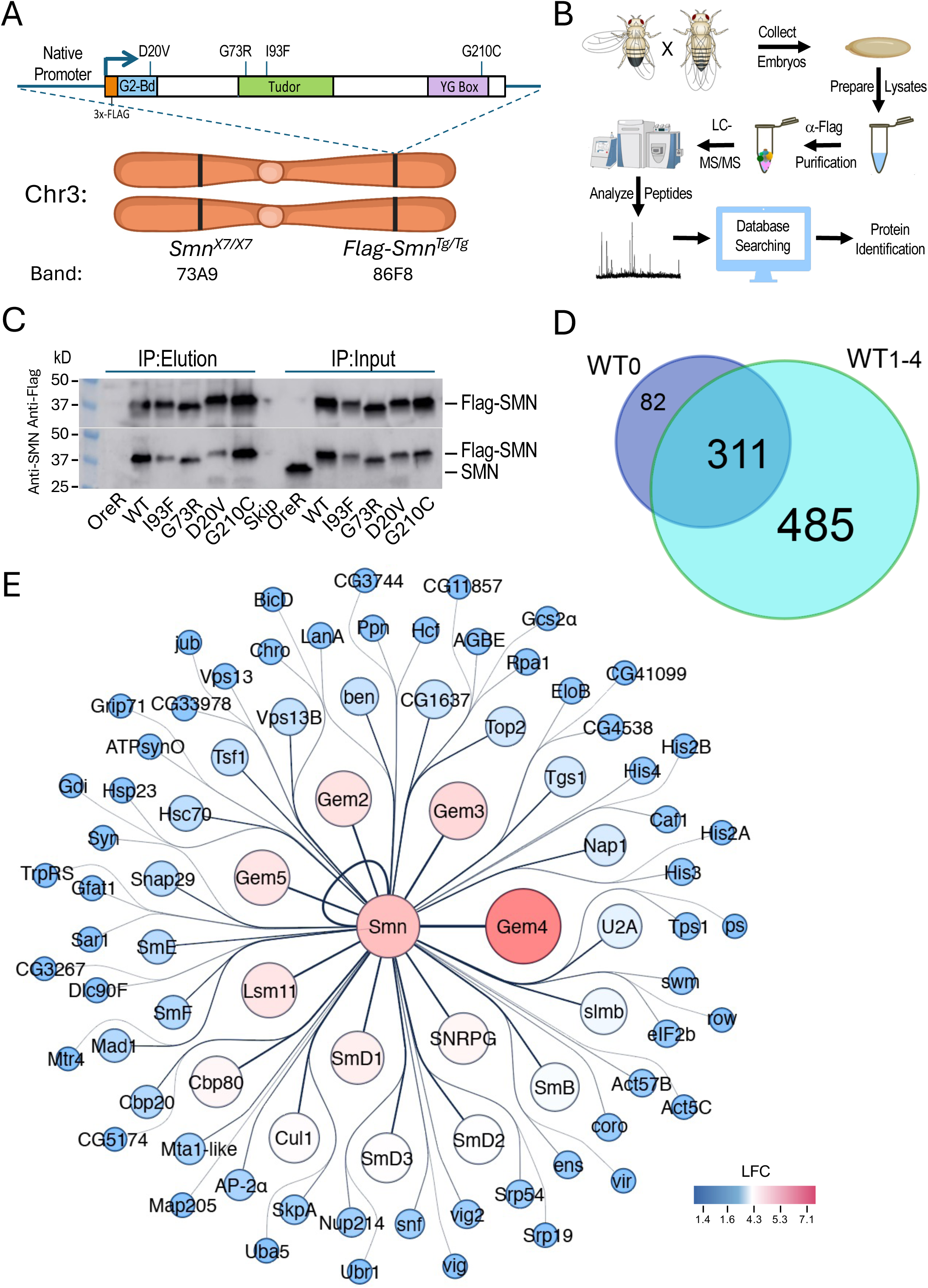
Experimental setup, workflow and overview of results. A) Ideogram of fruitfly chromosome 3 (Chr3) showing cytogenetic band locations of the endogenous *Smn* gene at 73A9 and the Flag-*Smn* rescue transgene (Tg) at PhiC31 landing site located at 86F8. Cartoon above shows the features of the rescue transgene driven by the native Smn promoter and control region. The SMN coding region was tagged with a 3x-FLAG epitope at the N-terminus. In addition to the WT *Smn* line, stocks expressing four different missense alleles (D20V, G73R, I93F and G210C) were also generated. B) Diagram of experimental workflow, beginning with embryo collection and ending with identification of peptides and protein predictions. C) Western blot of immunoprecipitation (IP) experiment for Oregon-R (OreR) control or the variou stable transgenic stocks described above. The upper blot was probed with anti-Flag and the lower blot was probed with anti-SMN, verifying the presence of the untagged endogenous protein in the OreR input lane but not in any of the other lanes. D) Venn diagram of the total number of proteins identified in Flag-IPs from *Smn^WT^* transgenic animals, comparing the four replicates generated here WT1-4 with that of a previous dataset, WT0 [34]. E) Graphical heat representation of the top protein hits as determined by Log2 fold-change (LFC) from the WT samples vs the OreR controls. Heatscale is at bottom right.

As outlined in Fig. 1B, we used population cages of these ‘proteomic stocks’ to set up embryo collections (0-24h) from which we prepared lysates. The embryonic lysates were then subjected to anti-Flag purification followed by LC-MS/MS, peptide mapping and protein identification (see Methods for details). Western blot analysis of the five Flag*-Smn* lines, along with a wildtype negative control (OreR) is shown in Fig. 1C, demonstrating the presence of endogenous (untagged) SMN only in the OreR input lane, and its absence from the immunoprecipitated material in all of the lanes.

### AP-MS analysis of wildtype SMN binding partners

Altogether, we performed three different mass spectrometry runs that included a total of 23 samples and identified a total of 894 different *Drosophila* proteins (Fig. 1D). Among the four WT biological replicates (WT1-4), there were 796 proteins that copurified with Flag-SMN (Fig. 1D), 79% of which (311/393) overlapped with those we identified previously (WT0) using a single replicate [34]. A visual summary of the top hits, as ranked by Log2 fold-change differences between WT and OreR controls, is shown in Fig. 1E. Clearly identified among the highest ranked partner proteins are the core members of the SMN complex (SMN, Gem2, Gem3, Gem4/Glos and Gem5/rig) and its best known RNP assembly clients, the Sm proteins (SmB, SmD1, SmD2, SmD3, SmE, SmF, SNRPG/SmG and Lsm11). Other RNP biogenesis factors include Cbp80•Cbp20, Snf, U2A and Tgs1. Also prominent atop this list are components of the SCF^slmb^ E3 ligase (Slmb, Cul1 and SkpA) and the E2 ubiquitin conjugase, ben (Fig. 1E) both of which have been previously reported to interact physically and genetically with SMN [34, 37]. These data show that many orthologs of the well known mammalian SMN binding partners co-purified with wildtype *Drosophila* SMN.

To assess the reproducibility of these interactions and to help establish appropriate cutoffs for downstream comparisons, we performed a Student’s T-test (two tailed, homoscedastic) between the WT and OreR replicates (four each) and generated a volcano plot versus the fold-change data (Log2 transformed). As shown in Fig. 2A, many of the aforementioned SMN partner proteins are well above the traditional threshold of p<0.05 (-Log10 > 1.25). Conspicuously below that line are other known SMN binding partners like SkpA, ben, Tgs1, Cbp20, SmD3 and Gem2, most of which are relatively small proteins (15-25 kDa). Gem2 and SMN form the heterodimeric core of the SMN complex [27, 51]; whereas Gem4 and Gem3 are not part of the core, but are much larger in size (100-120 kDa). Small and/or hydrophobic proteins have fewer peptides that can be detected by MS and thus are at a disadvantage compared to larger ones.

**Figure 2:**
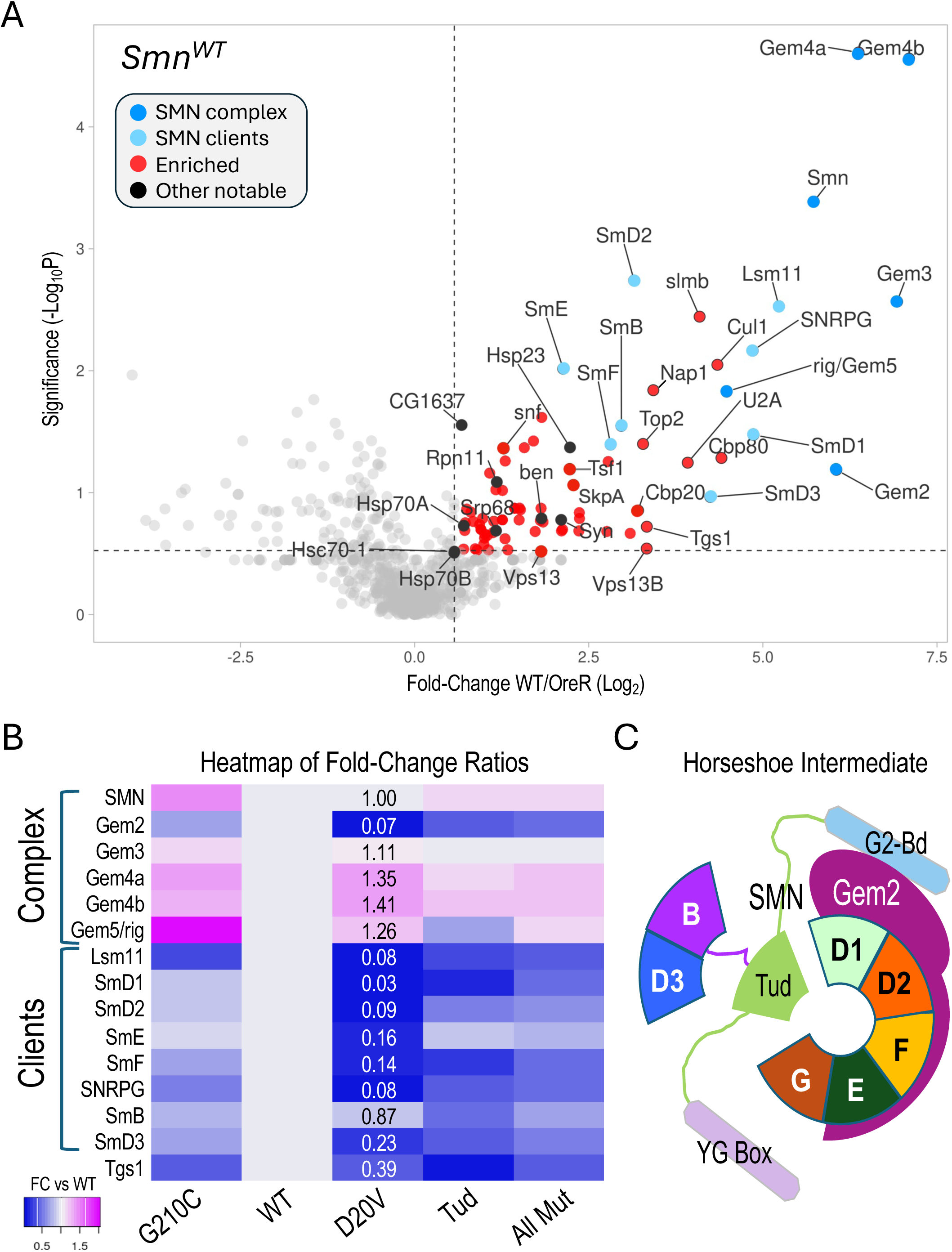
Analysis of AP-MS data. A) Volcano plot of the full *Smn^WT^* dataset. Dots represent individual proteins. The SMN complex is shown in dark blue, known SMN clients (Sm proteins) are shaded light blue. Other notable proteins are shaded black. Vertical dotted line represents a fold-change cutoff of 1.5x (LFC ≥ 0.58) for enriched proteins (shaded in red). Horizontal dotted line is shown for display purposes, see text for details regarding significance cutoffs. B) Heatmap of fold-change ratios for well-known SMN binding partners, comparing the data from the WT pulldowns to those of the G210C, D20V and Tud mutants. Tud = combined results for G73R and I93F. AllMut = combined results for all of the mutants. C) Cartoon of known intermediate in spliceosomal snRNP assembly pathway, showing the seven canonical Sm proteins (B, D3, D1, D2, F, E and G), Gemin2 (Gem2), and SMN (with its three differentially shaded domains corresponding to those in Fig. 1A). See text for details.

To determine if our MS data could be used to identify changes in the association of known binding partners among the SMA-causing missense alleles, we next focused on the members of the SMN complex and its well-known clients, the Sm proteins. For example, the human G279C allele (G210C in fly) is reported to be slightly hyper-oligomeric [27], whereas D44V (D20V in fly) is reported to reduce binding affinity to Gem2 [52]. The two Tudor domain (Tud) mutations (G73R and I93F) are known to cause temperature-sensitive misfolding [36, 53] and are plausibly expected to interfere with binding to Sm client proteins. Therefore, we generated a heatmap of fold-change ratios for various SMN alleles versus that of the WT (Fig. 2B). As shown, the G210C mutant pulled down slightly higher levels of SMN and Gemins 3-5, consistent with its reported hyper-oligomeric nature. Concordantly, the D20V mutant co-purified considerably less Gem2 than did the WT construct, despite the fact that there were nearly identical levels of SMN detected in the two pulldowns (Fig. 2B).

During U-snRNP assembly, Gem2 is known to directly bind to five of the seven Sm proteins, forming a key horseshoe-shaped intermediate (Fig. 2C). Interestingly, those same five proteins (SmD1, D2, E, F, G) were the most reduced in the D20V mutant pulldowns (Fig. 2B), whereas SmB and D3 were the two least affected clients. As predicted, the G73R and I93F mutants (Tud) also pulled down fewer Sm client proteins, but the contrast among the five Gem2-binders was less apparent (Fig. 2B). Moreover, association of the Sm clients was relatively unaffected in the G210C mutants Fig. 2B). Note that Lsm11 (which is also reduced in D20V) dimerizes with Lsm10 to replace the SmD1•D2 dimer in the context of the U7 snRNP [17]. Fold-change ratios for the combined set of mutants (AllMut) vs WT are provided for general comparison, as are data for the snRNP biogenesis factor, Tgs1. In summary, these data show that the binding profiles of the SMN missense mutants are consistent with their known activities in RNP assembly.

### Gene ontology profiling of wildtype and mutant SMN binding partners

The overall profiles of polypeptides identified by AP-MS in the wildtype and mutant Flag-SMN pulldowns were relatively similar (Fig. S1A), particularly among the top hits. Given the relatively mild phenotypes of these SMA models, this was perhaps unsurprising. Among the co-purifying proteins with fold-change values ≥ 1.5, the WT, G210C and D20V constructs pulled down 210, 239 and 223 partners, respectively (Fig. S1A). By contrast, the Tud mutants (G73R and I93F) copurified only 144 such proteins. This general trend of reduced binding to targets shared between WT and Tud mutants is illustrated in Fig. S1B, by a difference plot. A similar comparison of shared hits between WT and G210C, for example, showed a more or less equal distribution of increases and decreases (Fig. S1B). To gain insight into biological processes that might be most affected by these mutations, we carried out gene ontology/functional enrichment analysis using gProfiler [54] using the top 100 hits (measured by fold-change) from each of the SMN^WT^ and SMN^Tud^ experiments as queries. As shown in Fig. 3A, spliceosomal RNP assembly and mRNA processing categories (along with many overlapping ones not illustrated) were clearly the top hits in both samples.

**Figure 3:**
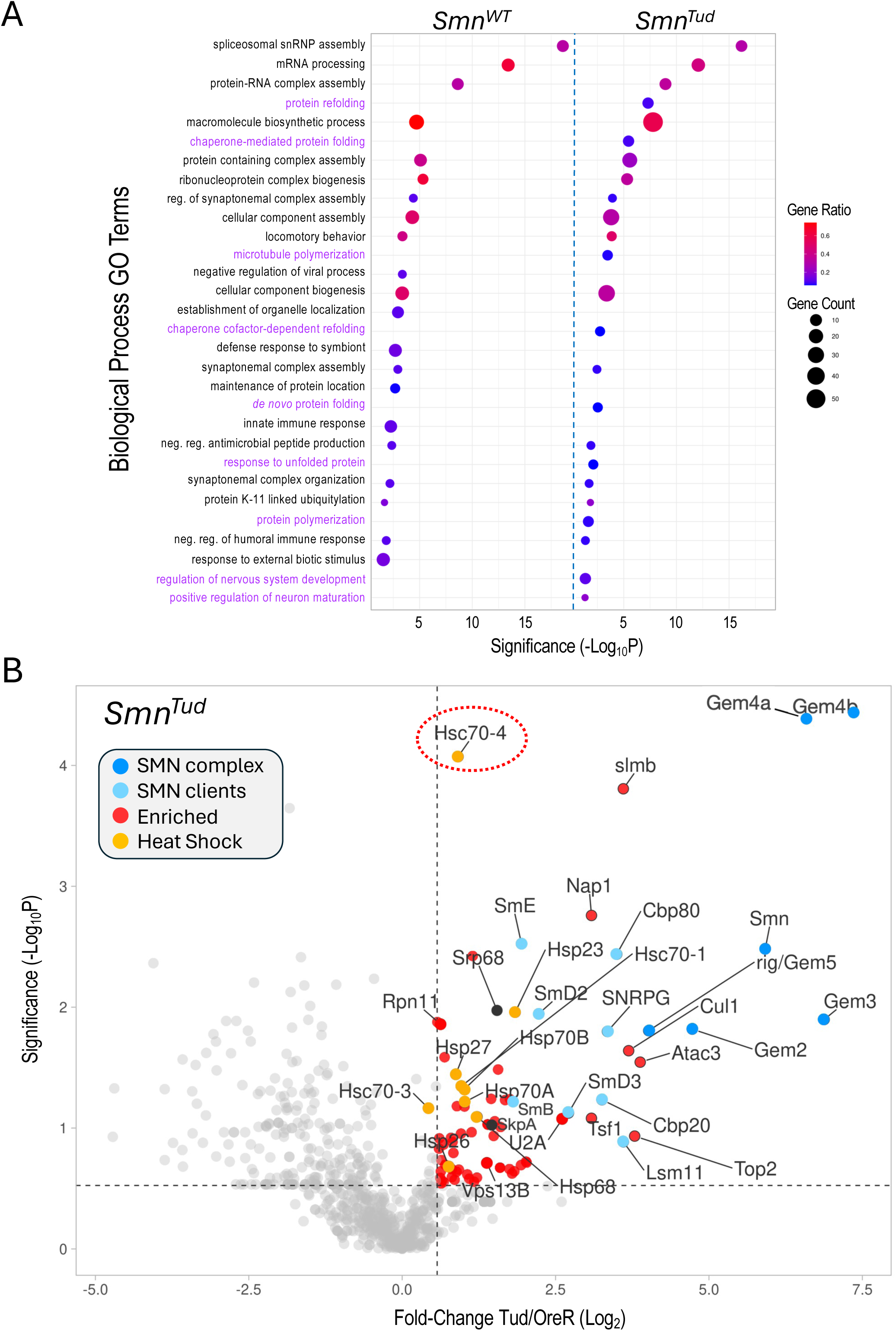
Comparative gene ontology (GO) analysis of the top 100 proteins identified in the *Smn^WT^* and *Smn^Tud^* AP-MS datasets. For each GO term listed, the size of the dot is proportional to the number of genes contained within that term (gene count), and the fraction of those genes scoring significantly (gene ratio) is represented using a heatmap (legend at right). Adjusted P-values (−log10 transformed) for each GO term were calculated and plotted separately for the WT and Tud results. B) A) Volcano plot of the full *Smn^Tud^* dataset. Dots represent individual proteins, shaded as per Fig. 2A and shown in the color key (inset). Heat shock proteins are highlighted in orange. Hsc70-4 is circled. See text for details.

Other shared categories include the regulation of synaptonemal complex formation, cellular component assembly and biogenesis, antimicrobial peptide synthesis, locomotory behavior, and the humoral immune response. Given that we and others have identified connections between SMN and many of these pathways including innate immune signaling [20, 34, 37, 55, 56], these categories were not unexpected. Prominent among the significant GO terms not shared by the two lists were those centered on aspects of protein folding and re-folding (Fig. 3A). Because the GO terms were chosen on the basis of fold-change data alone, we next wanted to assess the significance of the full set of proteins identified in the SMN^Tud^ pulldowns. We therefore generated a volcano plot of the Tud mutants compared to OreR controls. Many of the SMN^WT^ binding partners, including members of the SMN complex, the Sm clients, the Cbp80•20 cap-binding complex, the histone chaperone Nap1, and the SCF^slmb^ E3 ligase complex (Cul1, SkpA, slmb) were significantly enriched in the SMN^Tud^ pulldowns as well (Fig. 3B). The small HspB family member, Hsp23, was also significantly enriched in both the WT and Tud samples (Figs. 2A and 3B).

Conspicuous among the factors identified in the Tud pulldowns (Fig. 3B), are members of the large HspA family (ref) that includes Hsp70- and Hsc70-type proteins. Although many members of this family were also identified in the WT samples, they were not as highly enriched. Most prominent among the HspA family members that were significantly enriched in the SMN^Tud^ pulldown is Hsc70-4 (Fig. 3B, circled). This protein is particularly noteworthy because it has been recently linked to neuromuscular disease phenotypes in mouse and fruitfly models of both SMA and ALS (discussed below; [38, 39]. Furthermore, Hsc70 (subtype not specified) was originally shown to co-purify with SMN in HeLa cells [57], though its importance to SMA was unrecognized at the time (discussed below).

Because heat shock proteins constitute a major class of molecular (folding) chaperones, they interact with a large fraction of the proteome [6]. Thus HSPs are often identified as ‘contaminants’ in many AP-MS experiments. Nevertheless, the data in Fig. 3 identify clear signatures of a proteotoxic stress response (e.g. to unfolded proteins).

### Proximity labeling of human stress granule components, GEMIN3 and STRAP

In human cells, the SMN complex is known to localize to stress granules (SGs), non-membrane bound cytoplasmic structures that form in response to a variety of cellular stressors [58, 59]. Low levels of SMN are thought to impair the cell’s ability to form SGs [60]. Previous studies have used proximity labeling approaches (e.g. BioID, APEX) to analyze the composition of RNP granules, finding that many of the interactions that take place during SG assembly are pre-existing in unstressed cells [61–63]. One such study generated more than a hundred different stable cell lines expressing biotin ligase (BirA) tagged versions of RNP granule associated proteins, including two members of the human SMN complex, GEMIN3/DDX20 and STRAP/UNRIP [61]. As expected, these two proteins were among the top five BirA-tagged ‘baits’ to identify SMN as one of its biotinylated ‘prey’ (Fig. 4).

**Figure 4:**
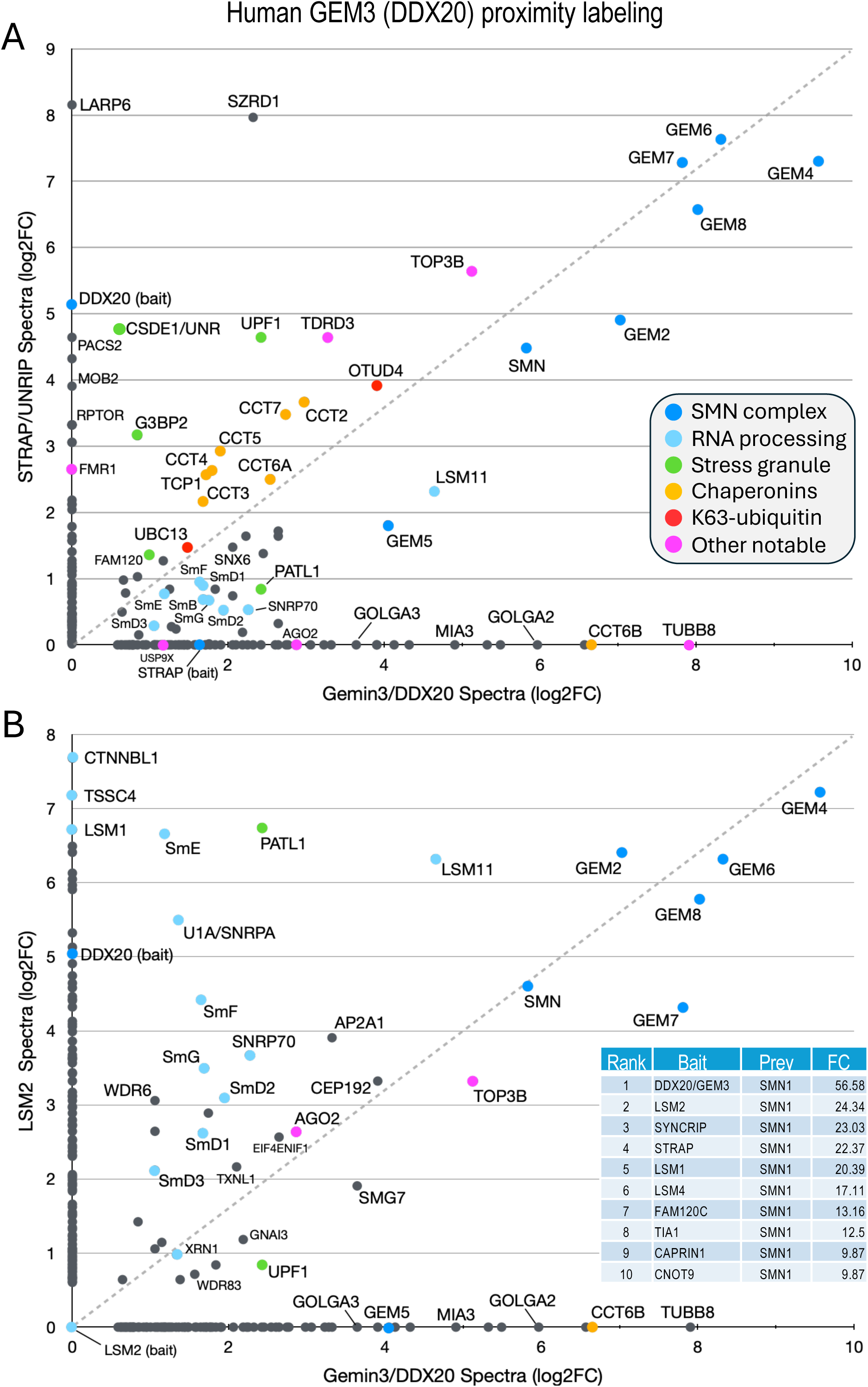
Comparison of previously published [61] BioID proximity labeling data for human stress granule proteins. Fold-change scatter plots of GEMIN3/DDX20 vs STRAP/UNRIP (A) and vs LSM2 (B) are shown. Color key is inset in panel A. The inset in panel B provides a table listing the the top ten BirA-tagged baits listing SMN as a prey are shown for comparison. See text.

Both STRAP and GEMIN3 (GEM3) are known to form complexes outside of their interaction with SMN [64–68]. STRAP (serine/threonine kinase receptor associated protein), originally identified as a component of the TGF-beta signaling machinery, is also known to interact with UNR/CSDE1 and LARP6 [65], serving as a translational regulatory factor[69]. STRAP is a peripheral member of the SMN complex, tethered via its direct interaction with GEMIN7 as part of the GEMIN6•7•8 subunit [70]. GEM3/DDX20 is a putative RNA helicase that heterodimerizes with GEMIN4 and has many reported activities [67, 68, 71-73]. To identify additional candidate factors and complexes that associate with the human SMN complex, we utilized the mass spectrometry data for proximity labeling of STRAP and GEM3 in unstressed cells [61] to generate a scatter plot (Fig. 4A). We reasoned that proteins forming associations with the SMN complex would tend to plot along the diagonal, whereas those that are uniquely associated with either GEMIN3 or STRAP would localize along the X or Y axes, respectively. Consistent with that interpretation, and with what is known about the interactions among and between members of the SMN complex [70], GEM4 and the GEM6•7•8 subunit are very strongly labeled, localizing to the upper right hand corner of the plot (Fig. 4A).

Other members of the complex, including GEM2, SMN and GEM5 align along the diagonal with the Sm client proteins. Of particular note, LARP6 and CSDE1 were well labeled only by STRAP, whereas the Golgin proteins GOLGA2 and GOLGA3 and the microtubule protein TUBB8 were detected only in the GEM3/DDX20 experiment (Fig. 4A). Proteins that are known to be important for stress granule formation (PATL1, FAM120, UPF1 and G3BP2) were identified by both baits but are not well aligned along the diagonal. In contrast, TDRD3 and TOP3B form a complex with FMRP that is implicated in synapse formation and neurological development [74, 75] and are labeled by both GEM3 and STRAP. Two factors involved in aspects of K63-linked ubiquitylation, UBC13 (bendless in flies) and OTUD4, and all eight members of the chaperonin ring complex (TCP1/CCT1 thru CCT8) were also labeled by both baits (Fig. 4A).

As a control, we plotted the proximity labeling data for one of the other top bait proteins for which SMN was identified as a prey, LSM2 (Fig. 4B, inset). LSM2 is an Sm-like protein with manifold connections to the mRNA surveillance machinery and is also part of the U6 snRNP particle, though it is not a known SMN client. As shown in Fig. 4B, factors involved in RNA processing and quality control (e.g. XRN1, UPF1, LSM1, TSSC4 and CTNNBL1) are well labeled by LSM2 but neither UBC13 nor the chaperonins were identified. These latter two categories are important because they point to a role for the SMN complex specifically in innate immune signaling and chaperone-mediated protein folding, both of which processes are deeply integrated with the proteostasis network (discussed below).

The fruitfly ortholog of STRAP is called wing morphogenesis defect (Wmd) and its structure is well conserved. Whether or not it forms part of the SMN complex in the fly is an open question. The Gem6•7•8 subunit is entirely missing from Drosophilid genomes, although these proteins are conserved in other insects [22]. Consistent with the absence of its potential Gem7 tether [70], neither Wmd nor any putative Gem6-8 paralogs were detected in our Flag-SMN pulldowns. We therefore conclude that Wmd is not part of the *Drosophila* SMN complex (Fig. 5).

**Figure 5:**
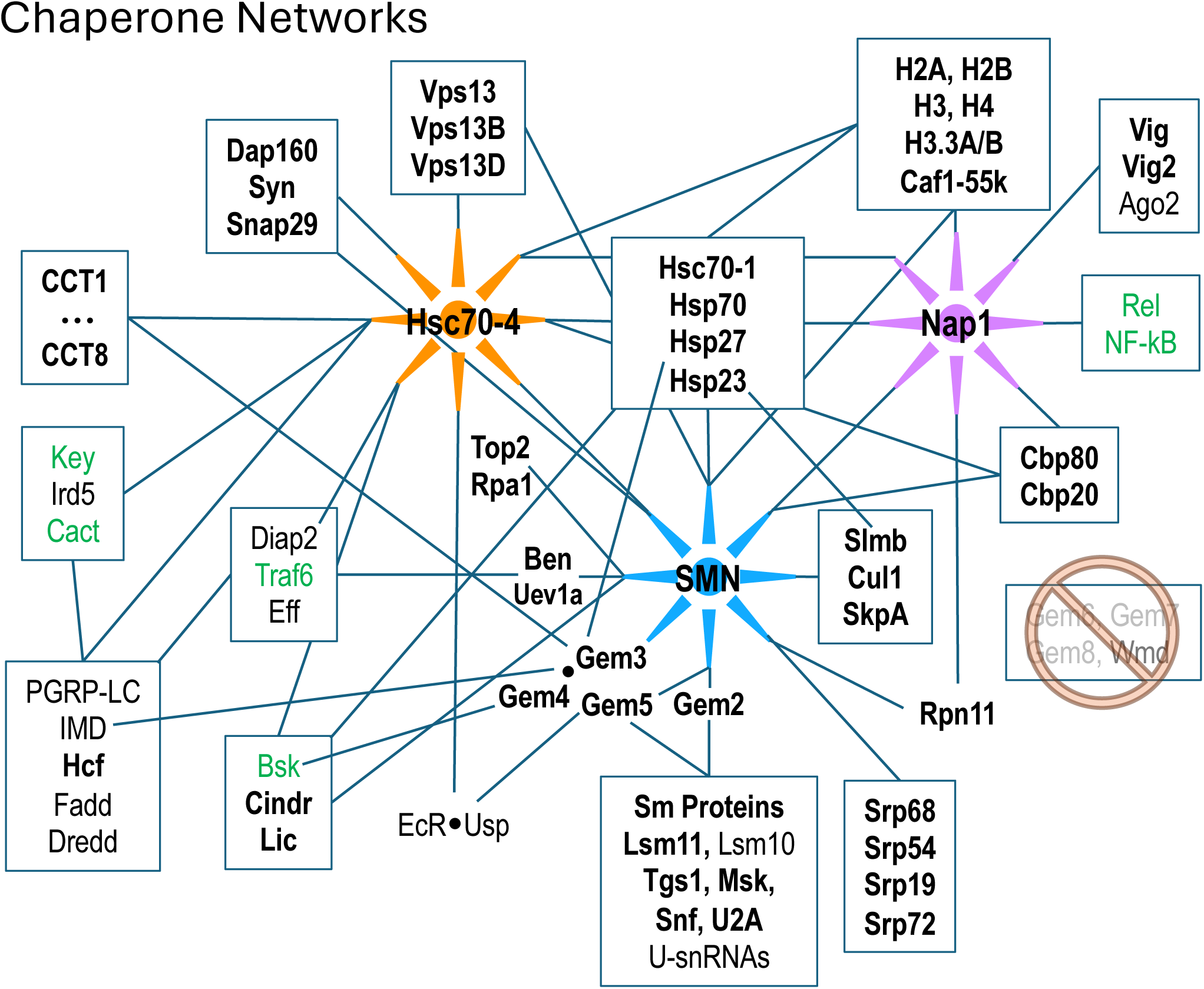
Diagram of protein-protein interactions between three key molecular chaperone systems: SMN (RNP assembly), Hsc70-4 (protein folding) and Nap1 (nucleosome assembly). Solid lines indicated known interactions. Factors listed in bold text are were identified in this work; those in green are innate immune signaling factors that were shown to interact genetically with SMN [37].

### Cellular components identified by purification of the Drosophila SMN complex

In addition to the many novel findings reported above, the AP-MS data confirm and extend our knowledge of cellular components and pathways that are connected to the SMN complex in diverse organisms. Beyond the obvious core members of the SMN complex and its Sm protein clients, these additional partners include, but are not limited to: the nuclear cap biding complex, the signal recognition particle, autophagic lumenal markers of the endoplasmic reticulum and Golgi, various kinds of ATPases, innate immune signaling factors such as kinases and ubiquitylases, synaptonemal complexes and other presynaptic cystosolic components, spindle proteins, chromatin remodelers (Table 1 and Fig. 5).

**Table 1:** Functional enrichment analysis of AP-MS data from. **Fig. 2A. GO terms focusing on cellular components are shown.**

| GO: Cellular Component | Term ID | P.adj | Term Size | Query Size | Intersection |
| --- | --- | --- | --- | --- | --- |
| SMN-Sm protein complex | GO:0034719 | 1.65E-25 | 11 | 12 | 9 |
| SMN complex | GO:0032797 | 1.12E-16 | 8 | 9 | 6 |
| intracellular anatomical structure | GO:0005622 | 2.41E-13 | 7266 | 203 | 174 |
| cytoplasmic U-snRNP body | GO:0071254 | 2.52E-13 | 7 | 9 | 5 |
| small nuclear RNP complex | GO:0030532 | 5.19E-12 | 60 | 32 | 9 |
| RNP complex | GO:1990904 | 3.79E-11 | 585 | 23 | 14 |
| membrane bounded organelle | GO:0043227 | 6.07E-08 | 5771 | 176 | 125 |
| non-membrane bounded organelle | GO:0043228 | 2.94E-07 | 1935 | 203 | 67 |
| U12-type spliceosomal complex | GO:0005689 | 3.3362E-05 | 12 | 12 | 3 |
| nBAF complex | GO:0071565 | 6.3162E-05 | 6 | 147 | 4 |
| SWI/SNF superfamily-type complex | GO:0070603 | 8.3337E-05 | 54 | 156 | 8 |
| U2-type spliceosomal complex | GO:0005684 | 8.4835E-05 | 35 | 22 | 4 |
| membrane-enclosed lumen | GO:0031974 | 0.00014107 | 1163 | 198 | 42 |
| ATPase complex | GO:1904949 | 0.00016633 | 118 | 170 | 11 |
| brahma complex | GO:0035060 | 0.00019953 | 16 | 147 | 5 |
| Cajal body | GO:0015030 | 0.00038333 | 14 | 22 | 3 |
| gamma-tubulin complex | GO:0000930 | 0.00119366 | 14 | 109 | 4 |
| SWI/SNF complex | GO:0016514 | 0.00132495 | 11 | 147 | 4 |
| spindle | GO:0005819 | 0.00137762 | 148 | 168 | 11 |
| nuclear cap binding complex | GO:0005846 | 0.001427 | 3 | 19 | 2 |
| nuclear body | GO:0016604 | 0.00176798 | 93 | 6 | 3 |
| presynaptic cytosol | GO:0099523 | 0.00285133 | 4 | 19 | 2 |
| U4/U6 x U5 tri-snRNP complex | GO:0046540 | 0.00323491 | 19 | 32 | 3 |
| signal recognition particle | GO:0048500 | 0.00371319 | 7 | 99 | 3 |
| presynapse | GO:0098793 | 0.00407557 | 239 | 139 | 12 |
| gamma-tubulin ring complex | GO:0000931 | 0.0049575 | 7 | 109 | 3 |
| nuclear lumen | GO:0031981 | 0.00614285 | 908 | 198 | 32 |
| UBC13-UEV1A complex | GO:0035370 | 0.01600461 | 2 | 101 | 2 |
| spindle midzone | GO:0051233 | 0.01732144 | 33 | 168 | 5 |
| CHD-type complex | GO:0090545 | 0.02295456 | 8 | 156 | 3 |
| pole plasm | GO:0045495 | 0.02777789 | 56 | 22 | 3 |
| pericentriolar material | GO:0000242 | 0.03015982 | 13 | 100 | 3 |
| cytoskeleton | GO:0005856 | 0.03018142 | 656 | 183 | 23 |

A few of these conserved connections deserve particular note. Perhaps unsurprisingly we identified numerous proteins with strong connections to RNP biogenesis and transport. U-snRNP components like U2A, Snf (U2B” and U1A’) and Lsm11•10 (U7 snRNP), the nuclear import factor Msk (Importin-7), the m7G-cap binding complex (Cbp80•20) and the trimethylguanosine synthase protein, Tgs1 (Table 1 and Fig. 5). Tgs1 was recently shown to play a role in snRNA 3’-end processing [76] with loss-of-function impacts in eye development but no obvious neuromuscular defects [55].

SMN has also been implicated in the biogenesis and/or regulation of other RNPs besides the U-snRNPs. For example, assembly of the signal recognition particle takes place in the nucleolus and cytoplasm, and this process is thought to involve the activity of the SMN complex [25]. In total, our AP-MS data identified Srp-19, -54, -68 and -72, suggesting that this putative function of the SMN complex is conserved in metazoa. Other connections to the nucleolus are evident; SMN and other Cajal body components are known to relocalize to the nucleolus during genotoxic stress [77, 78]. In support of these findings, we identified the large subunit of RNA pol I (Rpa1), topoisomerase 2 (Top2) and fibrillarin (Fib) in our dataset (Fig. 5). SMN, via its interaction with symmetric dimethylarginine (sDMA) residues on the pol II large subunit, is reportedly involved in R-loop resolution in the nucleoplasm [79, 80]. We hypothesize that this same type of activity could be important for RNA pol I transcripts in the nucleolus upon cell stress.

### RNP assembly chaperones and innate immunity

We recently showed that hypomorphic mutation or depletion of SMN induces a systemic hyperactivation of the Toll and IMD signaling pathways leading to formation of larval melanotic nodules [37]. Importantly, depletion of U-snRNPs due to mutation of *Phax* (an snRNA export factor) does not elicit this immune response [33]. Thus immune dysfunction in these SMA models is a direct consequence of reduced SMN levels and not a downstream or indirect consquence of snRNP loss [37]. As outlined in Fig. 5, there are numerous physical associations ([34, 81, 82]; this work) and genetic interactions [37] between members of the SMN complex and innate immune signaling factors like Ben, Traf6 and Imd. Not to be overlooked are conserved interactions between the small (B-type) heat shock ‘holdases’ like Hsp23 and Hsp27 and the SMN complex (Fig. 5). Interestingly, missense mutations in the human orthologues of two closely related small heat shock proteins, HspB8 (Hsp22) and HspB1 (Hsp27), are known to form abnormally strong interactions with GEM3 that are associated with motor neuron diseases [83]. Hence, our work points to the heterodimeric Gem3•Gem4 subunit of the SMN complex as a conduit between the RNP assembly chaperones, the folding chaperones/co-chaperones (heat shock proteins and chaperonins), and the innate immune system (Fig. 5).

As mentioned above, Wmd/STRAP forms a connection to the TGF-beta signaling pathway in mammals, but this link appears to be severed in flies due to loss of the Gem7 bridge. Perhaps a new signaling link was established in flies via the activity of Gem5/rig. This protein was originally thought to serve as a nuclear hormone receptor/co-factor [84], as it interacts physically and functionally with the ecdysone receptor, EcR. Subsequently, it was identified as the orthologue of human GEMIN5 [22, 26]. Together with its binding partner Usp, EcR forms interactions with Hsc70-4 and numerous other shared interactors [82, 85]. Ecdysone is the central driver of developmental decision making in arthropods (Ecdysozoa); indeed, pulses of similar steroid hormones are known to control the timing of developmental transitions in all types of animals [86].

Viewed in that light, it is perhaps unsurprising that there would be a link between the snRNP assembly machinery and regulators of cellular proliferation [87]. In animals, high levels of SMN protein are really only required when building an organism. In flies, SMN is maternally provided and its levels remain high throughout embryogenesis, dropping to basal levels during the three larval stages. It rises again during pupation (metamorphosis), only to fall back again to basal levels upon eclosion as adults [36, 88]. In mice, depletion of SMN protein later in development has relatively little consequence compared to early depletion [89]. Thus, high levels of SMN are required in rapidly proliferating cells. How do cells communicate these needs and coordinate them with other biosynthetic pathways across and between cellular compartments?

### Chromatin assembly chaperones and innate immunity

Given SMN’s primary cellular location in the cytoplasm, it was somewhat surprising to see so many highly significant GO terms (Table 1) for nucleosome remodeling complexes like SWI/SNF CHD, Brahma and nBAF. As mentioned above, connections between SMN and Cajal bodies or nucleoli are to be expected, but direct links to bulk chromatin are hard to fathom. It is important to remember that, like any other protein, nucleosomal subunits are born in the cytoplasm. Similar to Sm protein subcomplexes, histones are assembled into heterodimers prior to their nuclear import and incorporation into chromatin. The key finding in our AP-MS datasets is nucleosome assembly protein 1, Nap1 (Fig. 5). This novel, high-confidence SMN partner was reproducibly detected in both the wildtype and mutant Flag-pulldowns (Figs. 1E, 2A, 3B, S1B). Nap1 was also highly enriched in our previous study (WT0, Fig. 1D; [34]) and its co-purification with Flag-SMN was confirmed by western blotting [90].

Nap1 has well-established functions in chromatin remodelling and gene expression regulation [91, 92], but it is also thought to play a role in the innate immune response, particularly to viral infection [93, 94]. In humans, there are five Nap1-like genes (*NAP1L1-L5*), three of which are retrogenes that are expressed exclusively in the nervous system [95, 96]. *NAP1L1* and *NAP1L4* are the most ancestral family members, and depletion of NAP1L1 interferes with the nuclear translocation of RelA, the 65kD subunit of NF-kB, leading to a weakened TLR3 (Toll-like receptor 3) response [97]. Nap1L1 also interacts with A- and C-type (Hsc70 and Hsp90, respectively) heat shock proteins, which are known to interact with H3•H4 heterodimers in the cytoplasm [98, 99]. As illustrated in Fig. 5, Nap1 and SMN are central members of two different assembly chaperone systems that share conserved connections to the folding chaperones and the innate immune signaling system. From a conceptual standpoint, the findings reported here expand our understanding and appreciation for the extent to which macromolecular assembly chaperones intersect with innate immune signaling proteins within the larger context of the proteostasis and ribostasis networks.

### Chaperoning the chaperones: Heat shock proteins and motor neuron disease

As protectors of the proteome (and the RNPome), molecular chaperones of the HspA (70 kD) family are important players in intracellular signaling pathways because they regulate the folding and activity of signaling proteins [100]. Importantly, expression of a misfolded protein is known to change the profile of HSP binding partners [101]. We recently identified a subset of SMA-causing missense mutations in the Tudor domain of SMN (G73R, I93F, V72G and F70S) that are temperature-sensitive [36]. In response to relatively small increases in culturing temperature (e.g. from 25°C to 27° to 29°C), the Tud mutants display reduced SMN protein levels and fairly dramatic changes in organismal viability [36]. As detailed in Fig. 3, the SMN^Tud^ mutants exhibit striking changes in binding-partner profiles even at the ‘sub-clinical’ temperature of 25°C. Most notable among the many HSPs that co-purify with the Tud mutants is Hsc70-4. Reciprocally, a high throughput screen for Hsc70-4 (HspA8 in mammals) client substrates identified GEM3 and SMN as potential targets [101].

Most important, a G470R missense mutation in Hsc70-4/HspA8 was recently identified as a potent suppressor of the SMA phenotype in a murine disease model [39]. Monani and colleagues identified a spontaneous point mutation in the background of their mouse colony that suppresses SMA-like phenotypes by *bypassing the need for high levels of full-length SMN* [39]. They mapped the mutation to the substrate recognition domain of Hspa8 and found that this allele binds less efficiently to SMN [39]. Although the precise molecular mechanism behind the genetic suppression is unclear, it requires the presence of a transgene that expresses the truncated SMNΔ7 isoform. The authors posit that the reduced affinity (partial loss-of-function) of HspA8^G470R^ for SMN allows the cell to ‘repurpose’ the folding chaperone away from SMN and onto other clients (e.g. SNAREs) that are important for neurotransmission [39]. Consistent with this view, we also found that aberrantly-formed SMN complexes can change the profile of Hsc70 binding partners ([39, 101]; this work).

If changing the overall balance of well- vs. poorly-folded clients can redirect HSPs to different pools of substrates, then what happens to the clients upon changes in the pools of HSPs? Perhaps equally important in this regard are the findings of Zarnescu and colleagues [38]. These authors found that a dominant-negative (gain-of-function), ALS-causing allele of TDP-43 aberrantly sequesters Hsc70-4/Hspa8 mRNA away from translating ribosomes in mouse and *Drosophila* disease models [38]. Recent evidence also suggests that, in addition to proteins, HSPs may also assist in the folding of certain non-coding RNAs [102–104]. Ancestrally, Sm and Sm-like proteins are known to function as RNA chaperones. Could SMN be involved in regulating localization and/or translation of HspA8 mRNA? Interestingly, we previously showed that the SMN client, SmD3, is involved in the transport and localization of *oskar* mRNA in the fruitfly ovary [105]. Moreover, we also identified TDP-43 mRNA as a target of Sm proteins in human cells via RIP-seq [23]. Additional studies will be needed in this area to precisely determine the etiological overlap between SMA and ALS.

However, given the well-known roles of HspA8 and its close relatives in synaptic vesicle recycling and micro-autophagy, the idea that chaperone-related dysfunction lies at the root of two of the most prominent motor neuron diseases is very appealing. Taken together with our findings, these two studies [38, 39] provide clear examples of how the lines between loss-of-function and gain-of-function muations can become blurry when viewed in the context of a large network. Cseremly and colleagues posited that chaperone-related immune dysfunction was an *emergent property* of a distorted proteostasis network [106]. They defined an emergent property as one that “can not be elucidated from the properties of any single network element;” rather, it emerges as a consequence of interactions within the whole network [106]. Indeed, this notion that chaperone deficiencies or polymorphisms, might distort signaling networks in unpredictable ways to induce disease states has turned out to be rather prescient.

Despite the fact that the primary therapeutic agents used to treat SMA are splice altering drugs [107], defects in pre-mRNA splicing do not cause SMA. The drugs alter the normal splicing pattern of *SMN2* to increase levels of full-length SMN. Previous work in our laboratory and others strongly suggests that the underlying cause of the disease is related to functions of SMN that are independent of its role in spliceosomal snRNP biogenesis. Animals bearing hypomorphic SMN point mutations that cause milder forms of SMA complete development and display normal snRNP levels, but still exhibit neuromuscular defects [33, 35]. As outlined here, manifold connections among and between molecular chaperones like HspA/Hsc70, SMN, and Nap1 with innate immune signaling pathways are conserved between insects and mammals. We therefore hold that the neuromuscular dysfunction in SMA and ALS is a direct consequence of perturbations in the proteostasis network.

## Materials and Methods

### Fly stocks and husbandry

Oregon-R was used as the wild-type control. The creation of transgenic *Smn* fly lines was previously described [31]. Generation of ‘proteomic stocks’ (i.e. stable fly lines expressing only transgenic Flag-SMN protein in the absence of endogenous SMN) was also described previously [34, 35]. Briefly, virgin females from the *Smn^X7^*/TM6B-GFP (null/balancer) line were crossed to *Smn^X7^,*Flag*-Smn^TG^/*TM6B-GFP males at 25°C, where TG represents the WT, G73R, I93F or G210C allele. To reduce stress from overpopulation and/or competition from heterozygous siblings, crosses were performed on molasses plates with yeast paste and GFP negative (*Smn^X7^,*Flag*-Smn^TG^/Smn^X7^*) larvae were sorted into vials containing standard molasses fly food during the second instar larval stage. Large numbers of these progeny were allowed to intercross in order to establish stable stocks. Following expansion, a stable population was formed and then maintained for 1-5 years (35-150 generations). Although Flag-*Smn* was initially hemizygous, this process allowed for meiotic recombination and selection to generate populations that are essentially homozygous (spot-checked by PCR) at the Flag-*Smn* locus at band 86F but remain null at the endogenous *Smn* locus at band 73A.

### Embryo collection, sample preparation, and Flag-immunoprecipitation

0-24h embryos were collected from Oregon-R control and the various Flag-SMN stocks, dechorionated, flash frozen, and stored at -80C prior to use. Roughly 75-100 ul of the packed dechorionized embryos were used for each replicate, which were resuspended on ice into 300 ul of Lysis Buffer containing 100mM potassium acetate, 30mM HEPES-KOH at pH 7.4, 2mM magnesium acetate, 0.5mM dithiothreitol (DTT) plus ½ tablet of protease inhibitor cocktail. Lysates were generated inside a 1.5mL centrifuge tube by crushing with a pestle and then centrifuged for 10 min at 4°C at 13,000 rpm in a microfuge. The soluble (middle) layer of the lysate was transferred to a new tube, and assayed (Bradford) for protein content.

30ul of anti-FLAG M2 agarose beads (Sigma) were resuspended and then pre-washed in 1 mL lysis buffer (by inverting 5 times and then centrifuging at 100xg for 1 min) a total of four times. After pre-washes, beads were resuspended in 100 ul lysis buffer. For each replicate, 5 mg of embryonic lysate were then added to the prepared beads and incubated for 2 h at 4°C with end-over-end rotation. After incubation, beads were spun down (4°C, 500xg, 1min) resuspended in 1 mL of Wash Buffer 1 (20mM Tris•HCl, pH 8.0, 100 mM KCl, 2 mM MgCl2, 10% glycerol, 0.05% Tween-20, 0.5 mM DTT and 0.2 mM PMSF) and washed 2x with (4°C, 100xg, 1min) centrifugation. This process was repeated 3x in Wash Buffer 2 (same as WB1 except 250 mM KCl was used) and then a final time in Wash Buffer 1 for a total of six washing steps.

Samples were eluted 3x into 50ul of Elution Buffer (20mM Tris•HCl, pH 8.0, 100 mM KCl, 10% glycerol, 0.5 mM DTT, 0.2 mM PMSF) containing 200 ug/mL of 3xFLAG peptide (Sigma). The first two elutions were combined and small portions were used for subsequent quality control steps (10% each for silver staining and western blotting) prior to sending for mass spectrometic analysis. Once checked, the immunoprecipitated samples were subjected to SDS-PAGE and stained with coomassie. Lanes (∼1cm) for each sample were excised and the proteins were reduced, alkylated, and in-gel digested with trypsin overnight at 37°C. Peptides were extracted, desalted with C18 spin columns (Pierce) and dried via vacuum centrifugation. Peptide samples were stored at -80°C until further analysis.

### Liquid chromatography coupled with tandem mass spectrometry (LC-MS/MS)

Immunoprecipitated samples were analyzed by LC-MS/MS using an Easy nLC 1200 coupled to a QExactive HF mass spectrometer (Thermo). Samples were injected onto an Easy Spray PepMap C18 column (75 µm id × 25 cm, 2 µm particle size; Thermo) and separated over a 90 min time period. The gradient for separation consisted of 5–45% mobile phase B at a 250 nl/min flow rate, where mobile phase A was 0.1% formic acid in water and mobile phase B consisted of 0.1% formic acid in 80% ACN. The QExactive HF was operated in data-dependent mode where the 15 most intense precursors were selected for subsequent fragmentation. Resolution for the precursor scan (m/z 350–1700) was set to 60,000, while MS/MS scans resolution was set to 15,000. The normalized collision energy was set to 27% for HCD. Peptide match was set to preferred, and precursors with unknown charge or a charge state of 1 and ≥ 7 were excluded.

### Data analysis

Raw data files were processed using Proteome Discoverer version 2.4 (Thermo Scientific). Peak lists were searched against a reviewed Uniprot drosophila database (downloaded Feb 2020, containing 21,973 sequences), appended with a common contaminants database, using Sequest HT within Proteome Discoverer. The following parameters were used to identify tryptic peptides for protein identification: 10 ppm precursor ion mass tolerance; 0.02 Da product ion mass tolerance; up to two missed trypsin cleavage sites; carbamidomethylation of Cys was set as a fixed modification; oxidation of Met and acetylation of N-terminus were set as variable modifications. Scaffold (version 4.7.3, Proteome Software) was used to validate MS/MS based peptide and protein identifications. Peptide identifications were accepted if they could be established at greater than 95% probability to achieve an FDR less than 0.1% by the Scaffold Local FDR algorithm. Protein identifications were accepted if they could be established at greater than 99.0% probability and contained at least 2 identified peptides. Relative quantitation was performed using the calculated quantitative values (spectral counts) within Scaffold. The data will be deposited to the ProteomeXchange Consortium via the PRIDE partner repository [108] using the dataset identifier PXDxxxxx.

Once identified by mass spectrometry, proteins were assigned to specific UniProt IDs. Official gene names, FBgn IDs, annotations, and a variety of other pertinent information used in this study were obtained from FlyBase [109]. Gene ontology (GO) analyses were performed with FBgn IDs using g:Profiler [110, 111]

## Acknowledgements

We thank C. Hamilton for assistance with fly husbandry and embryo collection, N. Barker-Krantz for sample preparation and MS data collection, B. McMichael and V. Vandadi for help with data visualization. This research is based in part upon work conducted using the UNC Proteomics Core Facility, which is supported in part by NCI Center Core Support Grant (P30-CA016086) to the UNC Lineberger Comprehensive Cancer Center. This work was supported by NIH/NIGMS grant R35-GM136435 (to AGM).

---

**Supplementary Fig S1:**
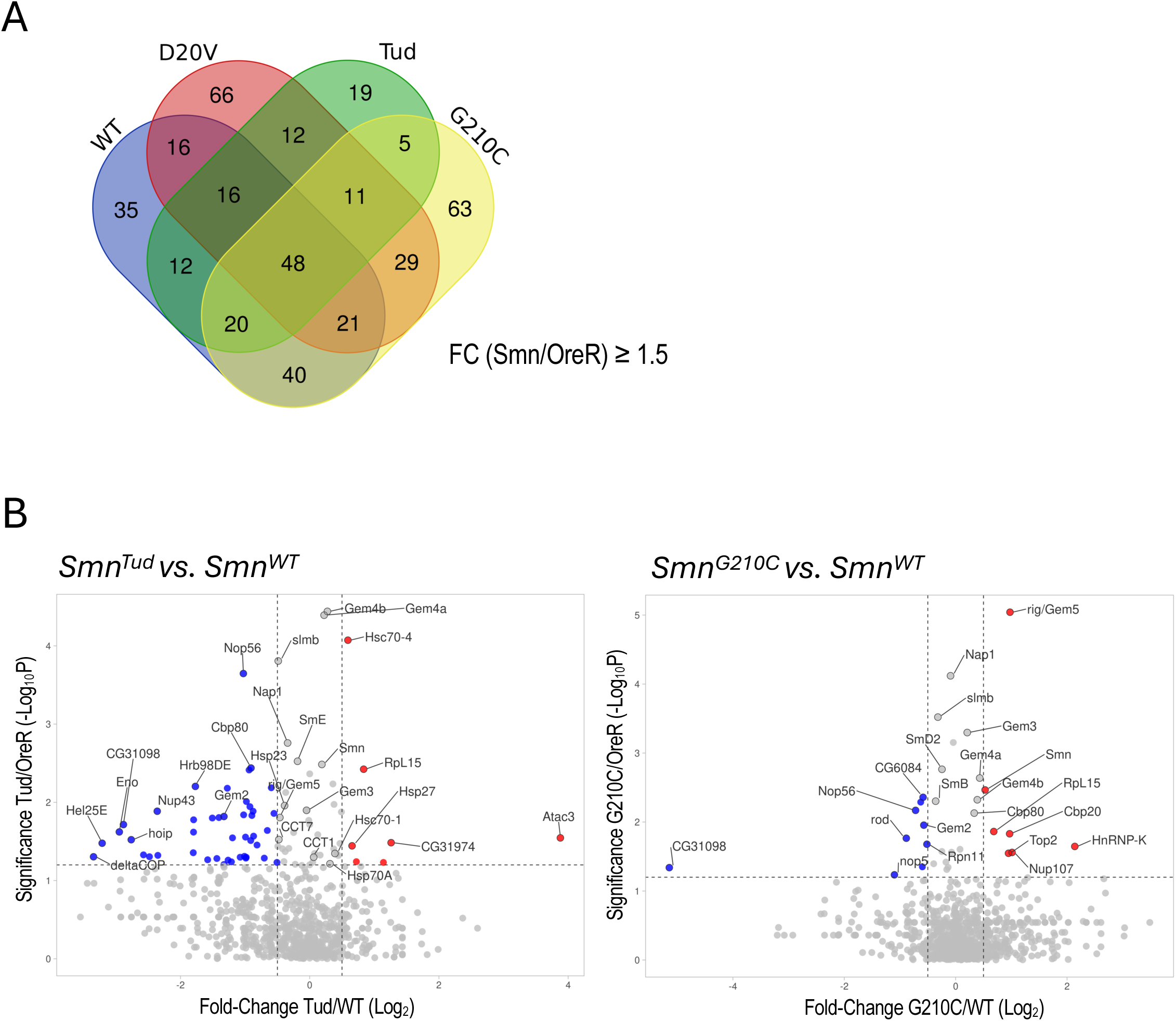
A) Venn diagram of proteins enriched (>1.5x) in AP-MS pulldowns of WT, D20V, Tud or G210C constructst. B) Difference scatter plots of shared proteins identified in the Tud vs WT or G210C vs WT AP-MS experiments.

